# 3D Printed Ti_3_C_2_T_x_ MXene/PCL Scaffolds for Guided Neuronal Growth and Photothermal Stimulation

**DOI:** 10.1101/2023.08.28.555002

**Authors:** Jianfeng Li, Payam Hashemi, Tianyi Liu, Ka My Dang, Michael G.K. Brunk, Xin Mu, Ali Shaygan Nia, Wesley D. Sacher, Xinliang Feng, Joyce K. S. Poon

**Author notes:** Corresponding authors at: Max Planck Institute of Microstructure Physics, Weinberg 2, Halle, 06120 Germany E-mail addresses (Jianfeng Li), (Joyce K. S. Poon).

## Abstract

The exploration of neural circuitry is essential for understanding the computational mechanisms and physiology of the brain. Despite significant advances in materials and fabrication techniques, controlling neuronal connectivity and response in three dimensions continues to present a formidable challenge. Here, we present a method for engineering the growth of three-dimensional (3D) neural circuits with the capability for optical stimulation. We fabricated bioactive interfaces by melt electrospinning writing (MEW) of 3D printed polycaprolactone (PCL) scaffolds followed by coating with titanium carbide (Ti_3_C_2_T_x_ MXene). Beyond enhancing hydrophilicity, cell adhesion, and electrical conductivity, the Ti_3_C_2_T_x_ MXene coating enabled optocapacitance-based neuronal stimulation due to illumination-induced local temperature increases. This work presents a strategy for additive manufacturing of neural tissues with optical control for functional tissue engineering and neural circuit computation.

## 1. Introduction

Deciphering neuronal activity in the brain is a longstanding and formidable challenge, holding significant implications ranging from advancements in computer architecture to clinical breakthroughs [1]. Neural circuits and synaptic connectivity constitute the fundamental building blocks of neural functionality [2]. Understanding the intricacies of brain circuitry not only inspires ways to enhance computational efficiency [3], but also provides invaluable insights into neurological disorders and pharmaceutical screening [4]. However, investigations in neural circuits *in vivo* or *ex vivo* are subjected to regulatory constraints and the complexities of biological brain structures [5]. *In vitro* neural circuit models can serve as simplified platforms for investigating neural functions at the cellular level. Although *in vitro* two-dimensional (2D) neural circuit engineering can be achieved by manipulating the surfaces in cell cultures[6], this fails to fully recapitulate the three-dimensional (3D) characteristics of the *in vivo* neuronal microenvironment. Various techniques, including microfluidics [7], electrocompaction [8], photolithography [9], colloids [10], and superparamagnetic nanoparticles [11], have been employed for engineering 3D neural circuits, but these methods lack precise control over structural and cellular details, limiting their suitability for manipulating neural circuits at the single-cell level. 3D printing can also enable the engineering of 3D neural circuits, but current 3D-printed neural circuit scaffolds either exhibit large millimeter-scale feature sizes [12] or require complex synthetic modifications of biological cues [13].

The precise modulation of neuronal activity within a circuit is critical to the comprehensive investigation of neural circuit function [14]. Optogenetics and multielectrode arrays (MEAs) are key technologies for neural stimulation and readout in cultures [15, 16]. However, optogenetics encounters challenges such as variable transfection efficiency and clinical approvals, while MEAs have limited spatial resolution in recording/stimulation and do not match the mechanical properties of tissues[17, 18]. The optocapacitive effect, the change in the cell membrane capacitance due to light-induced localized heating of the neurons, is an emerging approach for neuron stimulation without the need for genetic modifications and device transplantation [19]. Laser pulses can stimulate action potentials, but the optical energies and intensities must be kept sufficiently low to avoid damaging the tissue. The optical intensity for photodamage or phototoxicity in cells is substantially higher for red light (wavelength near 640 nm) than at blue, green, and ultraviolet wavelengths [20–22]. Wäldchen, S. et al.[20] show that intensity limit of photodamage is in excess of 10 MW/m^2^ for a wavelength of 640 nm, while Emon, B. et al.[21] recommends 57 W/m^2^ as a safe threshold for cell force homeostasis and Dubois, A. et al.[22] observed that 0.2 MW/m^2^ did not cause cellular damage in tissues. To stimulate with lower pulse energies and high spatial specificity, light absorbing particles, such as 3D fuzzy graphene [23], transition metal carbide/nitride (MXene) flakes [24], as well as gold or carbon nanoparticles [19], are incorporated onto the neurons to increase the photothermal conversion efficiency in localized regions of the cells. In this way, neurons can be stimulated at the μm and millisecond (ms) spatial and temporal scales. However, the application of these materials has been largely limited to 2D systems or single-cell studies, rather than to 3D structures.

In this work, we demonstrate Ti_3_C_2_T_x_ MXene-coated 3D polycaprolactone (PCL) scaffolds fabricated with melt electrospinning writing (MEW) with micrometer-scale features for optocapacitive neuron stimulation. Cell culture interfaces strongly influence cell properties such as cell adhesion, morphology, migration, and intercellular communication [25], thus bioactive interfaces can guide neuronal growth cones and the interconnection of neurons, enabling the controlled formation of neural circuits. Ti_3_C_2_T_x_ MXene is attractive for bioactive neural interfaces, since it is biocompatible and capable of strong photothermal effects due to its strong optical absorption of near-infrared wavelengths enhanced by the excitation of localized surface plasmons [26, 27]. Ti_3_C_2_T_x_ MXene has been used as a coating for cell culture substrates and as a medium suspension for dorsal root ganglion cell culture and stimulation [24]. However, these investigations have been limited to 2D systems and did not investigate neuronal interconnection. Additionally, Ti_3_C_2_T_x_ MXene coating can confer bioinert electrospun PCL conduits with excellent biocompatibility *in vitro* and *in vivo*, facilitating physiological electrical signal transmission and promoting angiogenesis [28]. Our Ti_3_C_2_T_x_ MXene-coated PCL scaffolds were effective in controlling neuronal interconnection, showing the potential to control neuronal network formation and activity.

## 2. Results and Discussion

### 2.1. 3D Fabrication of Ti_3_C_2_T_x_/PCL Scaffolds

**Fig. 1** shows an overview of the 3D Ti_3_C_2_T_x_/PCL scaffold neural interface. The Ti_3_C_2_T_x_ MXene serves as a surface modification for the 3D printed PCL scaffolds. Neuron growth is guided along the scaffold and the enables optical stimulation. PCL was selected as the substrate material because it is approved by the Food and Drug Administration (FDA) and has been explored extensively in tissue engineering for its ease of processing, biocompatibility, controllable biodegradation [29]. In the initial step, PCL scaffolds were fabricated through MEW-based 3D printing, achieving high-resolution structures with micrometer-scale features. This approach avoids the use of toxic solvents that may compromise the cytocompatibility of the supportive structures. Subsequently, pristine PCL scaffolds were coated with Ti_3_C_2_T_x_ MXene via drop casting and self-assembly, using a similar method as reported before [30]. The Ti_3_C_2_T_x_ MXene coating introduces extensive oxygen/fluorine-containing groups to the hydrophobic PCL surface, promoting cellular protrusion attachment [31]. Combined with the precisely defined microscale geometry, the 3D Ti_3_C_2_T_x_/PCL scaffold provides guidance for neurons to grow and form interconnected networks.

**Fig. 1:**
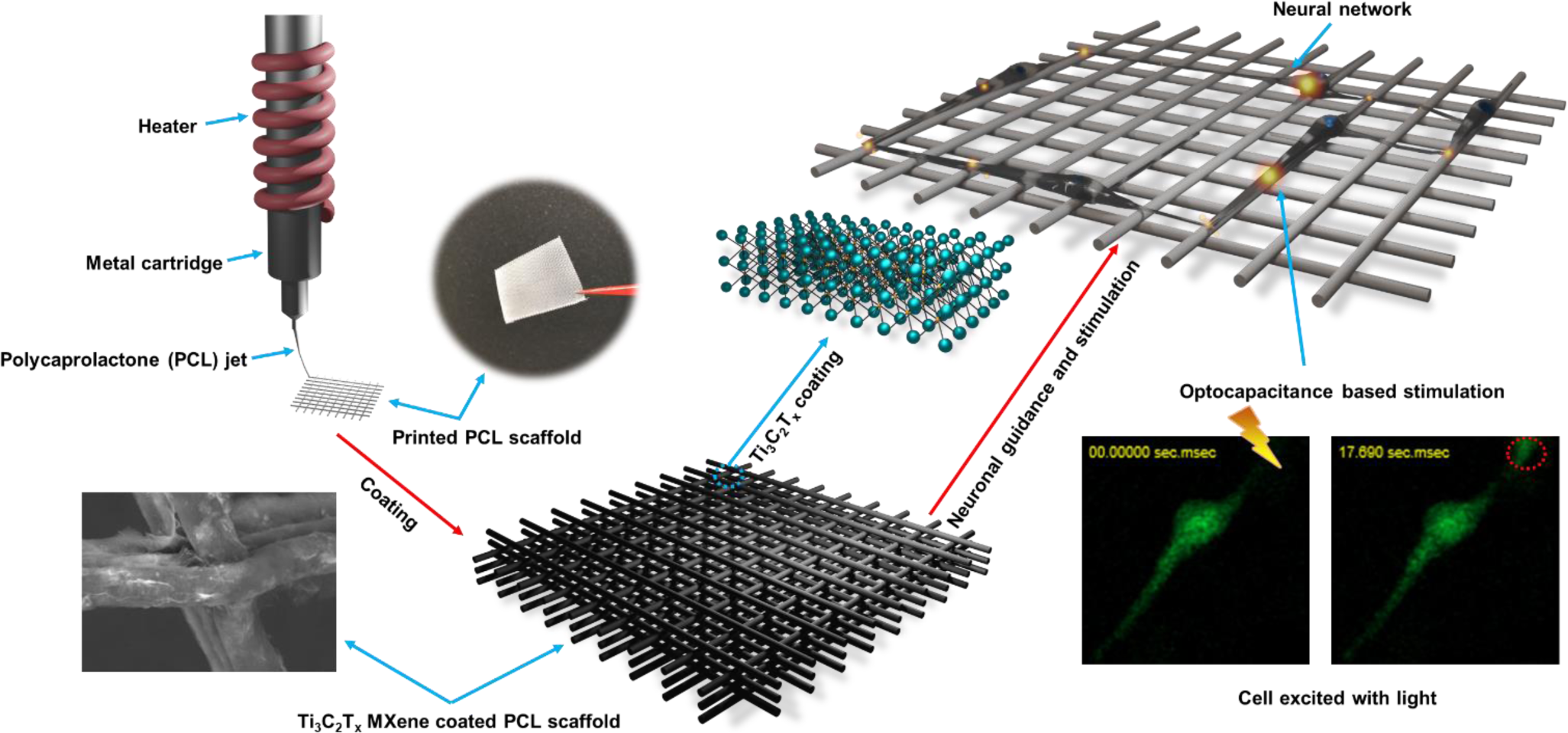
Schematic illustration of 3D fabrication of Ti_3_C_2_T_x_/PCL scaffold as neural network guidance and addressable optical stimulation platform. Mechanically robust PCL scaffolds were printed with the well-controlled PCL jet from the heated cartridge and Ti_3_C_2_T_x_ coating formed uniformly on the pristine PCL scaffold surface to render a cell friendly biointerface, which could be used to increase the effectiveness of optocapacitance based neuronal stimulation.

### 2.2. Characterization of Ti_3_C_2_T_x_ MXene and 3D Fabricated Scaffolds

To prepare Ti_3_C_2_T_x_ MXene, the Al layer in the precursor Ti_3_AlC_2_ MAX phase (**Fig. 2a**) was selectively etched away by acid, as described in the experimental section, to obtain the Ti_3_C_2_T_x_ MXene particles with the so-called accordion-like multilayer structures (**Fig. 2b**). With subsequent treatment, delaminated Ti_3_C_2_T_x_ MXene was obtained (**Fig. 2c**). Analysis of the X-ray diffraction (XRD) patterns revealed that the (002) peak, corresponding to the crystallographic plane of MXene, remained as the prominent peak following the etching, intercalation, and delamination treatment of the MAX phase (**Fig. 2d**). This observation suggests the successful removal of other components from the MAX phase. Furthermore, the broadening and shifting of the (002) peak from the MAX phase to MXene indicate a reduced thickness of the Ti_3_C_2_T_x_ MXene layer and an increased d-spacing [32]. The resulting aqueous dispersion of MXene was employed for the coating of the PCL scaffold in the next steps.

**Fig. 2.**
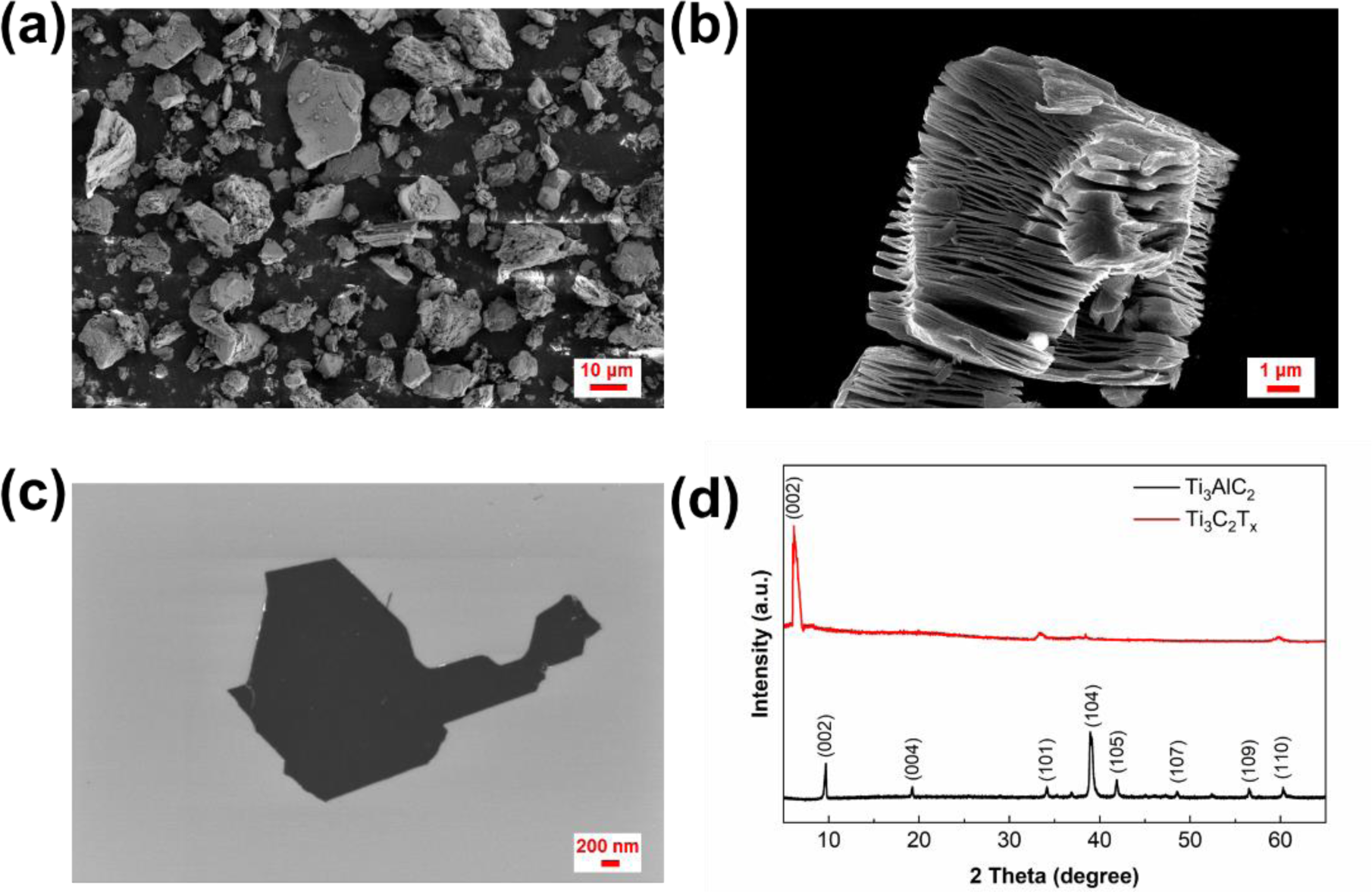
Ti_3_C_2_T_x_ MXene characterization. Scanning electron microscope (SEM) images of (a) Ti_3_AlC_2_ (MAX phase) powder particles (precursor of MXene), (b) Multilayer Ti_3_C_2_T_x_ (MXene particle) after etching away the Al layer, and (c) delaminated Ti_3_C_2_T_x_ MXene flake. X-ray diffraction (XRD) patterns of the (d) Ti_3_AlC_2_ MAX phase and Ti_3_C_2_T_x_ MXene.

Electrospinning is effective for fabricating one-dimensional (1D) and 2D nano/micro fibrous structures, with applications in numerous research domains, including biomedical engineering [33]. However, a challenge in electrospinning is the control over the generated fibers due to the chaotic motion of materials in the fabrication process [34]. To address this issue, MEW-based 3D printing was developed by placing the fluid jet onto the programmable motorized collector before the Plateau-Rayleigh instability criterion applies to the ejected fiber to attain accurate material deposition with high single fiber resolution [35]. With well-tuned MEW parameters, a 7-mm-thick printed construct can be formed with well-defined fiber placement and overall geometry [36]. Besides printing fine well-aligned straight fibers, MEW can also generate curved or coiled fibers when operating below the critical translation speed (CTS) and produce thinned fibers or beads above CTS [37]. MEW provides more control over the final geometry of the fibrous scaffolds than the conventional technology (i.e., depositing fibers without control and forming random pores) [37], enabling free infiltration of live cells through the structure. With the continual advancements, MEW can enable new possibilities for tissue engineering applications.

In our study, with precisely tuned parameters, 3D PCL scaffolds were fabricated by MEW with well-defined geometry and desirable morphology (**Fig. 3a; Supplementary Movie S1)**. Despite the delicate and fibrous nature of the printed structure, 3D printed PCL scaffold with more layers were sufficiently robust to be handled with tweezers (**Supplementary Fig. S1)**. Since PCL is hydrophobic [38], the 3D printed pristine PCL scaffolds with different strand distances (SDs) of 200 µm, 100 µm, and 50 µm (measured from the center of the strands) exhibited poor wettability of water with corresponding contact angles of 127.40 ± 4.12° (n = 3), 134.92 ± 2.02° (n = 3), and 133.36 ± 0.97° (n = 3), respectively (**Fig. 3a)**. However, a solution of 70% ethanol/water (70% EtOH; v/v) could spread throughout the structure, which eliminated the electrostatic attraction between the scaffold and the glass, causing the scaffold to detach from glass substrate (**Supplementary Movie S2**). Materials with low wettability, such as hydrophobic materials, exhibit limited cell adhesion and fail to provide the necessary environment for cellular growth and development [39]. To modify the wettability of PCL, we coated the structures with Ti_3_C_2_T_x_ MXene, which is hydrophilic due to abundant oxygen and fluorine functional groups on its surface.

**Fig. 3.**
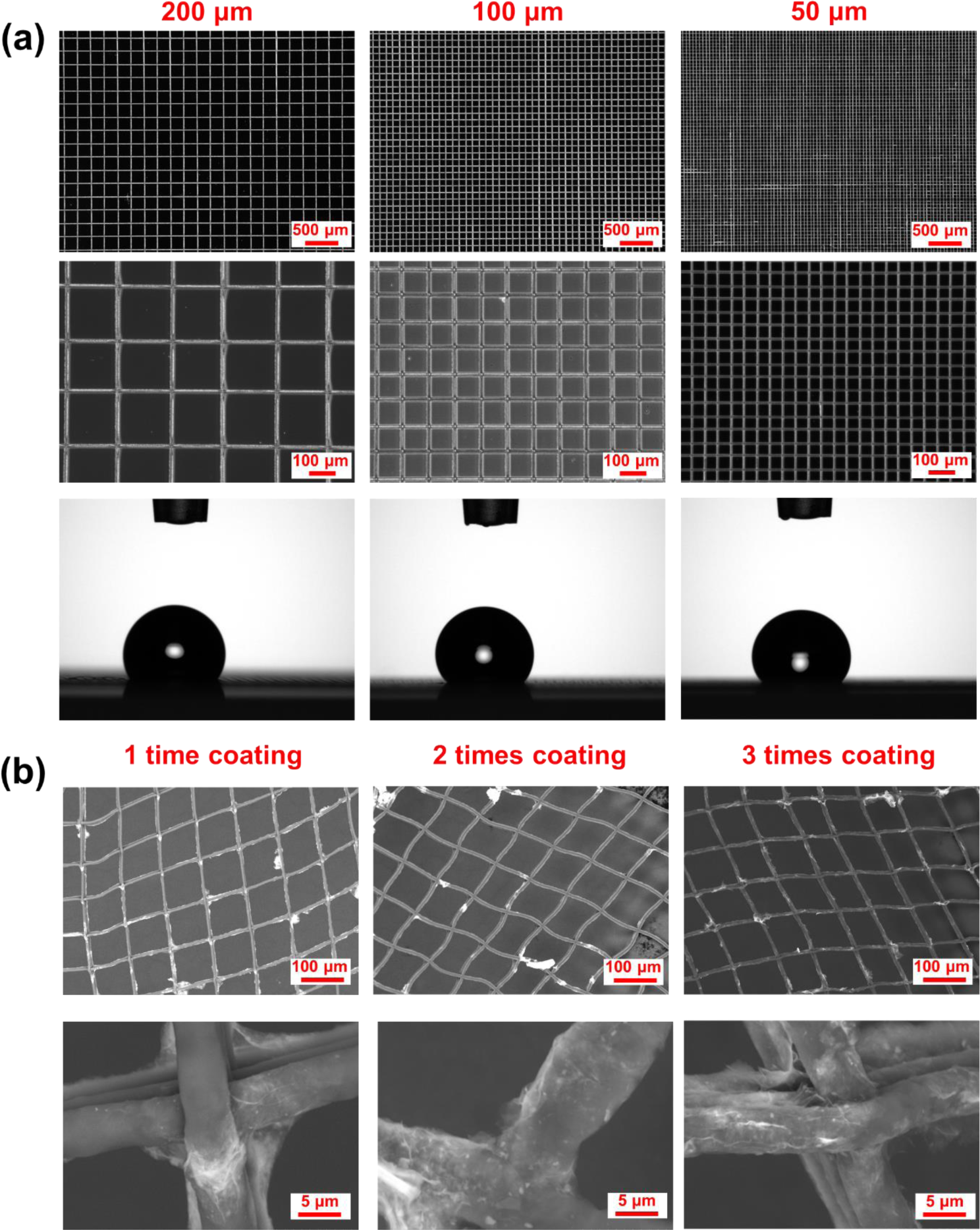
Characterization of 3D printed PCL and Ti_3_C_2_T_x_/PCL scaffolds. (a) Optical microscopy images of 3D-printed PCL scaffolds with different strand distances (SDs) (200 µm, 100 µm, and 50 µm) and corresponding water droplet profiles on the structures. (b) SEM images of 3D Ti_3_C_2_T_x_/PCL scaffolds with different coating times (1-3) at low and high magnifications.

We printed PCL scaffolds with random distribution of pore sizes ranging from 1 to 50 μm to investigate the impact of Ti_3_C_2_T_x_ MXene dispersion concentration and coating time on the resulting morphology of the coated structures. Ti_3_C_2_T_x_ MXene dispersions in 70 % EtOH, at concentrations of 0.5 mg/mL, 1.5 mg/mL and 3.0 mg/mL, were successfully and uniformly deposited onto the 3D printed PCL scaffolds with only minimal blockages observed in small pores (**Supplementary Fig. S2)**. However, when a highly concentrated Ti_3_C_2_T_x_ MXene dispersion (15.0 mg/mL) in water was used for coating, it resulted in the partial blockage of nearly all pores after a single coating, and subsequent coating led to the formation of a film covering the entire scaffold (**Supplementary Fig. S2**). This phenomenon can be attributed to the high viscosity of the concentrated dispersion and poor wettability during the coating process without the presence of ethanol. Furthermore, increasing the MXene concentration and the number of coating times led to lower electrical sheet resistance (**Supplementary Fig. S3**), as more interconnected conductive pathways were formed in the plane of the MXene structures [40]. For instance, the sheet resistance of 3-time coated PCL structure with 15.0 mg/mL MXene solution reached ∼ 18 Ω/, comparable to the 5 μm thick Ti_3_C_2_T_x_ MXene membrane reported in previous research [41]. The sheet resistance of the 3-time coated PCL structure with 3.0 mg/mL MXene dispersion was about 811 Ω/, and the structures preserved good morphology with minimal blockage of the pores (**Supplementary Fig. S2**). The electrical conductivity of these structures, along with the well-coated MXene structures on the 3D printed PCL scaffolds, are favourable properties for applications to cell cultures[42]. Additionally, we studied the wettability of Ti_3_C_2_T_x_ MXene coating on 3D printed PCL scaffolds with 100 µm SD. The Ti_3_C_2_T_x_ MXene coated PCL scaffolds showed favorable wettability (**Supplementary Movie 3**), with water droplet completely spreading into the coated structure within 1 min. The average diameter of deposited fiber was 5.75 ± 0.33 μm (n = 10), which is smaller than the diameter of neuronal cells, thereby allowing single cell attachment on the strands to facilitate the formation of precise cellular networks. When the MXene EtOH dispersion was applied to the pristine PCL scaffolds, the dispersion wetted the scaffolds easily, and the positively charged PCL surface attracted the negatively charged MXene flakes, resulting in uniform coatings on the scaffold [43, 44]. As shown in **Fig. 3b**, 3D MXene/PCL scaffolds exhibited good overall morphology without altering the original design of the substrate structure and MXene coatings showed improved coverage with an increasing number of coatings as shown in the zoomed in SEM images. This also explains the lower electrical resistance as the number of coating layers increases (**Supplementary Fig. S3**).

### 2.3. Neuronal Behaviour on the 3D Ti_3_C_2_T_x_/PCL Scaffold

Due to the low interfacial free energy and bio-inert surface chemistry of PCL scaffold, cells have limited adhesion on the structure, resulting in reduced controllability over tissue regeneration [45]. With the bioactive functional groups on the surface of PCL scaffold introduced by efficient Ti_3_C_2_T_x_ MXene coating, cell adhesion on the surface increased significantly after 1 day in culture, as shown in the fluorescence images in **Fig. 4**. Generally, more cells adhered and are distributed on the scaffold surface with Ti_3_C_2_T_x_ MXene coatings formed from higher concentrations of Ti_3_C_2_T_x_ MXene dispersions. Ti_3_C_2_T_x_/PCL scaffolds with 15.0 mg/mL Ti_3_C_2_T_x_ MXene dispersion coating had the densest cell population since the Ti_3_C_2_T_x_ MXene could block the pores and provide more adhering points for cells compared to scaffolds coated with diluted Ti_3_C_2_T_x_ MXene dispersions (**Supplementary Fig. S2**). Only a small number of cells adhered to the pristine PCL scaffold, and those cells showed rounded morphology and a lack of interaction with other cells through cilia, as shown in the SEM images in **Fig. 4**. In contrast, on the Ti_3_C_2_T_x_ MXene coated scaffolds, the cells showed elongated morphology with extended filopodia, spreading along the scaffolds. These results indicate that Ti_3_C_2_T_x_ MXene coating efficiently increased the bioactivity of a bioinert material surface and promoted synaptic connectivity.

**Fig. 4:**
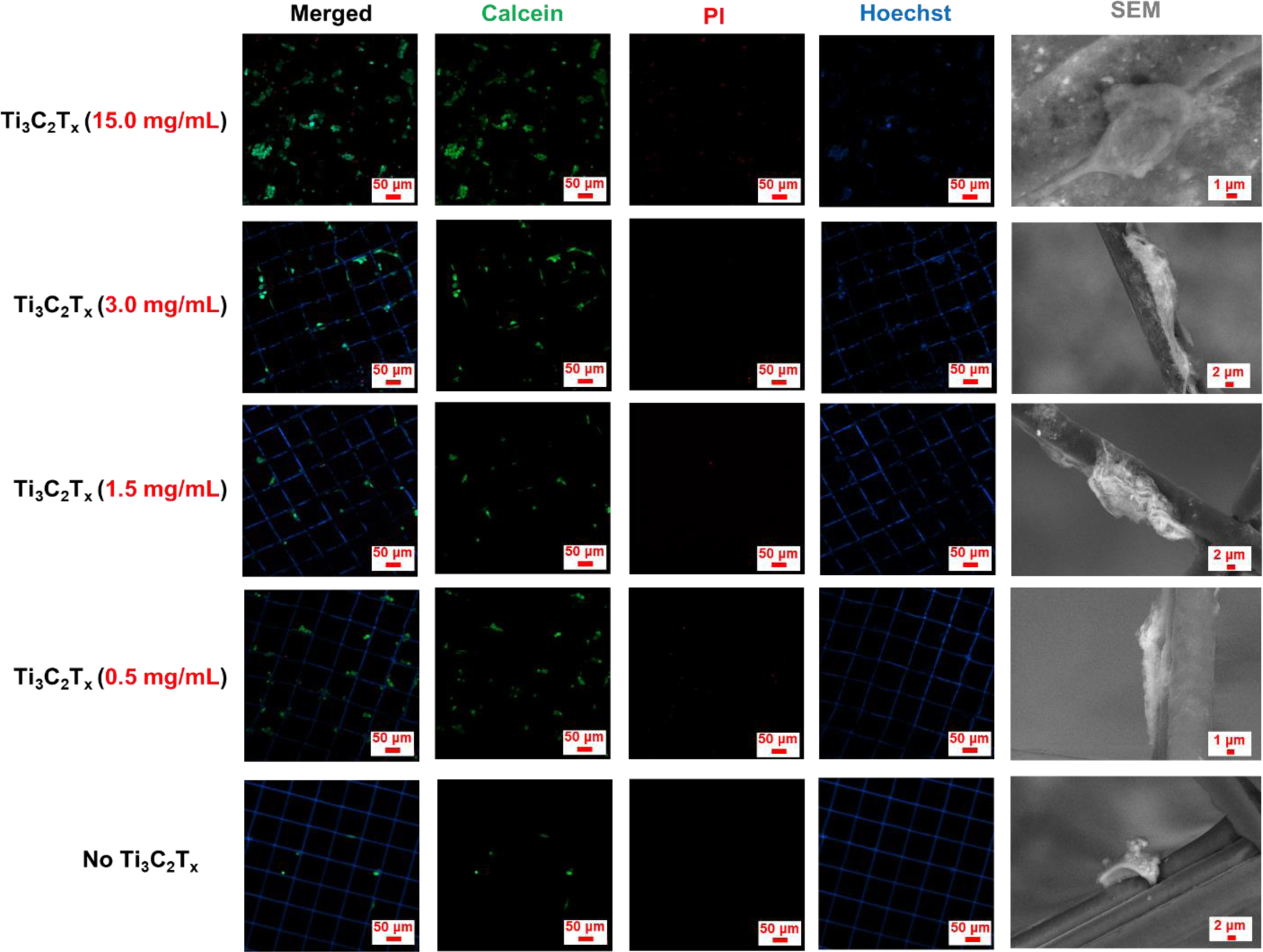
Cell viability and adhesion on 3D printed PCL scaffolds with and without Ti_3_C_2_T_x_ MXene coatings. Fluorescence images of cell adhesion and morphology on Ti_3_C_2_T_x_/PCL scaffolds with different coating concentrations of Ti_3_C_2_T_x_ MXene dispersions after 1 day of culture, followed by corresponding SEM images. Live cell (Calcein; green), dead cell [propidium iodide (PI); red] and cell nucleus with PCL substrate (Hoechst; blue).

**Fig. 5** shows the morphology analysis of the neurons on the scaffolds. The results again show that without the coating of Ti_3_C_2_T_x_ MXene, cells cultured on the pristine PCL scaffold exhibited a rounded morphology following seeding, whereas cells on Ti_3_C_2_T_x_/PCL demonstrated an elongated morphology with extended filopodia, as observed in the immunofluorescence image (**Fig. 5a**). Subsequent exposure to retinoic acid for 3 days resulted in the differentiation of cells on both substrates into dopaminergic lineage, characterized by elongated cell morphology (**Fig. 5b**). However, on the pristine PCL substrate, no interconnected neurons were observed, whereas multiple interconnected neurons grew along the Ti_3_C_2_T_x_/PCL, exhibiting significantly higher expression of F-actin and the neuronal marker β-tubulin III compared to cells on pristine PCL at Day 3, as well as both cultures at Day 1. As both F-actin and β-tubulin III are crucial proteins involved in neural growth signalling and navigation[46], the elevated expression of these proteins in cells on Ti_3_C_2_T_x_/PCL could be attributed to enhanced neuronal growth and interaction. The physicochemical interaction between cells and the Ti_3_C_2_T_x_ MXene coating also improved the interface for dopaminergic neuron differentiation and the formation of inter-neuronal circuits in our 3D structures.

**Fig. 5:**
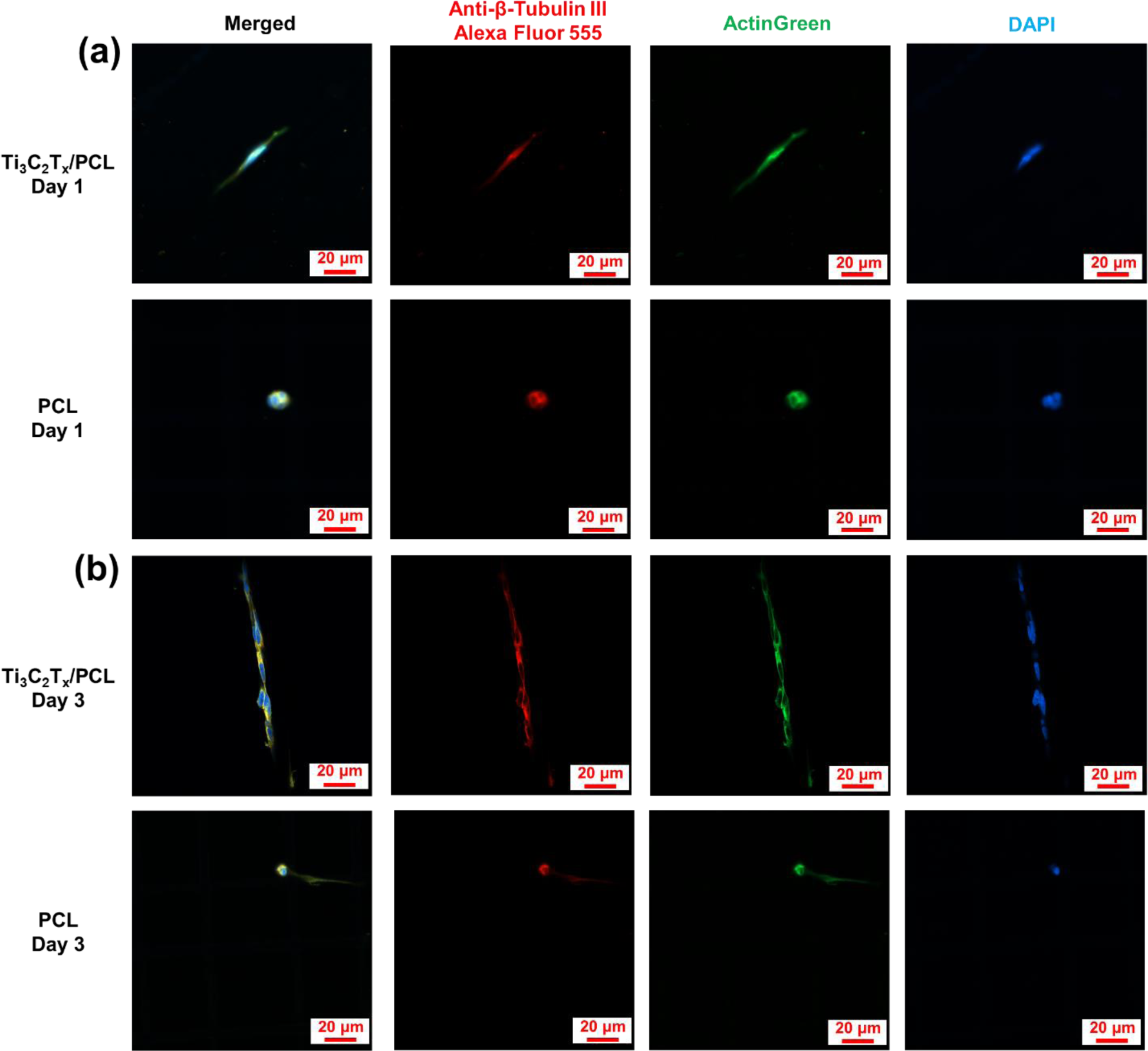
Neuronal morphology analysis on the Ti_3_C_2_T_x_ MXene (3.0 mg/mL dispersion) coated PCL and pristine PCL scaffolds. Fluorescence images of immunostained cultures towards dopaminergic lineage induction at (a) Day 1 and (b) Day 3, correspondingly. Neuronal marker β-tubulin III (Alexa Fluor 555; red), F-actin (AlexaFluor™488 phalloidin; green), and nuclei [4′,6-diamidino-2-phenylindole (DAPI); blue].

The cytocompatibility of the Ti_3_C_2_T_x_/PCL scaffolds was evaluated following a 7-day culture. As shown in **Fig. 6**, Ti_3_C_2_T_x_ MXene coating exhibited excellent cytocompatibility for neuronal cultures, while the SD of the printed scaffolds had an impact on neuronal growth and interconnection (**Fig. 6 and Supplementary Fig. S4**). More specifically, neurons did not strictly align along the scaffold strands during growth when the SD was smaller than 50 µm. This deviation from the anticipated growth pattern may have arisen from the interaction between growth cones of adjacent neurons when they were in close spatial proximity. In contrast, when the SD exceeded 100 µm, neurons adhered to the growth guidance provided by the scaffold strands, even as cellular multiplication occurred. This observation highlights the potential of Ti_3_C_2_T_x_/PCL scaffolds with larger SD for long-term applications in neuronal circuitry engineering, as they facilitate the desired alignment and interconnection of neurons.

**Fig. 6:**
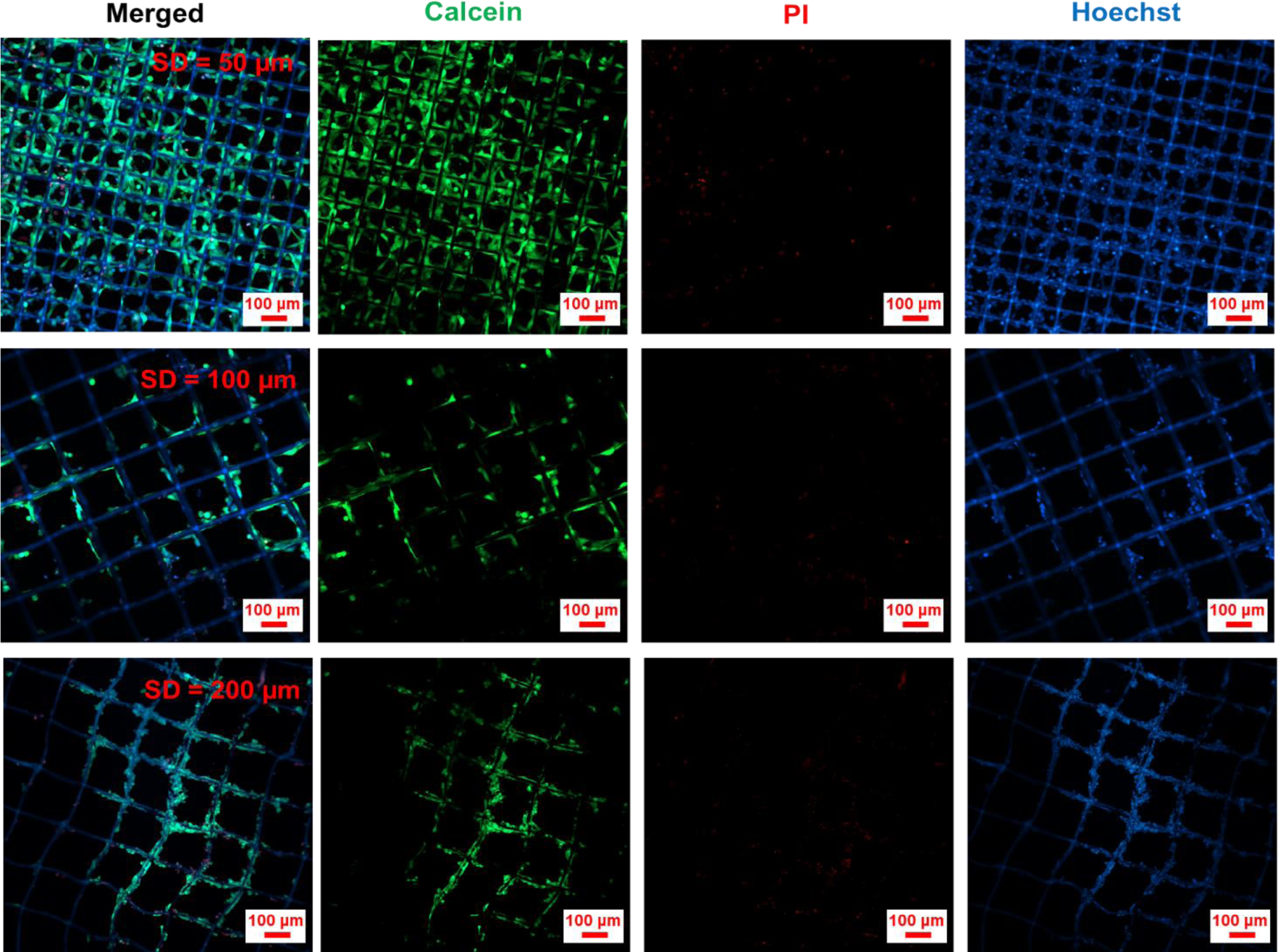
Cell viability and morphology on the 3D Ti_3_C_2_T_x_/PCL scaffolds with different SDs (50 µm, 100 µm, and 200 µm) after 7-day culture. Live cell (Calcein; green), dead cell [propidium iodide (PI); red] and cell nucleus with PCL substrate (Hoechst; blue).

Cell proliferation on the pristine 3D PCL scaffolds and Ti_3_C_2_T_x_/PCL scaffolds with varying concentrations of Ti_3_C_2_T_x_ MXene coatings was analyzed over 7 days by quantifying the area covered by live cell fluorescent staining. As shown in **Fig. 7**, the number of cells proliferating on Ti_3_C_2_T_x_ MXene-coated PCL scaffolds surpassed those on pristine PCL scaffolds consistently from Day 1 to Day 7, with a generally higher cell count associated with higher concentrations of Ti_3_C_2_T_x_ MXene coatings. In particular, the cell numbers on Ti_3_C_2_T_x_/PCL scaffolds coated with 3.0 mg/mL (Day 7) and 15.0 mg/mL (Day 1 and Day 7) Ti_3_C_2_T_x_ MXene dispersions were significantly higher than the cell numbers on pristine 3D PCL scaffolds (Day 1 and Day 7), with the cell count on the 15.0 mg/mL Ti_3_C_2_T_x_ MXene-coated scaffolds being 312 times higher than that on pristine PCL scaffolds (Day 7). The exceptionally high cell count observed can be attributed to the larger cell-supporting area created by the Ti_3_C_2_T_x_ MXene coating, which blocked pores in the scaffolds (**Supplementary Fig. S2**), while the loose adhesion of cells on the pristine PCL scaffolds may also cause cell loss during culture. Conversely, the poor adhesion of cells on pristine PCL scaffolds may have resulted in cell loss during the culture period. The lack of increase in cell numbers on 0.5 mg/mL Ti_3_C_2_T_x_ MXene-coated scaffolds from Day 1 to Day 7 can be attributed to the low coverage and stability of the coating derived from the highly diluted Ti_3_C_2_T_x_ MXene dispersion during cell culture. This limited growing space and caused detachment of cells from the substrate. Statistical analyses further reveal the significant impact of both Ti_3_C_2_T_x_ MXene coating concentration [*F*(4,20) = 142.65, *P* < 0.01] and culture time [*F*(1,20) = 116.34, *P* < 0.01] on cell proliferation. Additionally, the interaction between Ti_3_C_2_T_x_ MXene coating concentration and culture time was found to be significant [*F*(4,20) = 56.75, *P* < 0.01]. Hence, the Ti_3_C_2_T_x_ MXene coating exhibits the potential to provide an enhanced substrate for neuron proliferation.

**Fig. 7:**
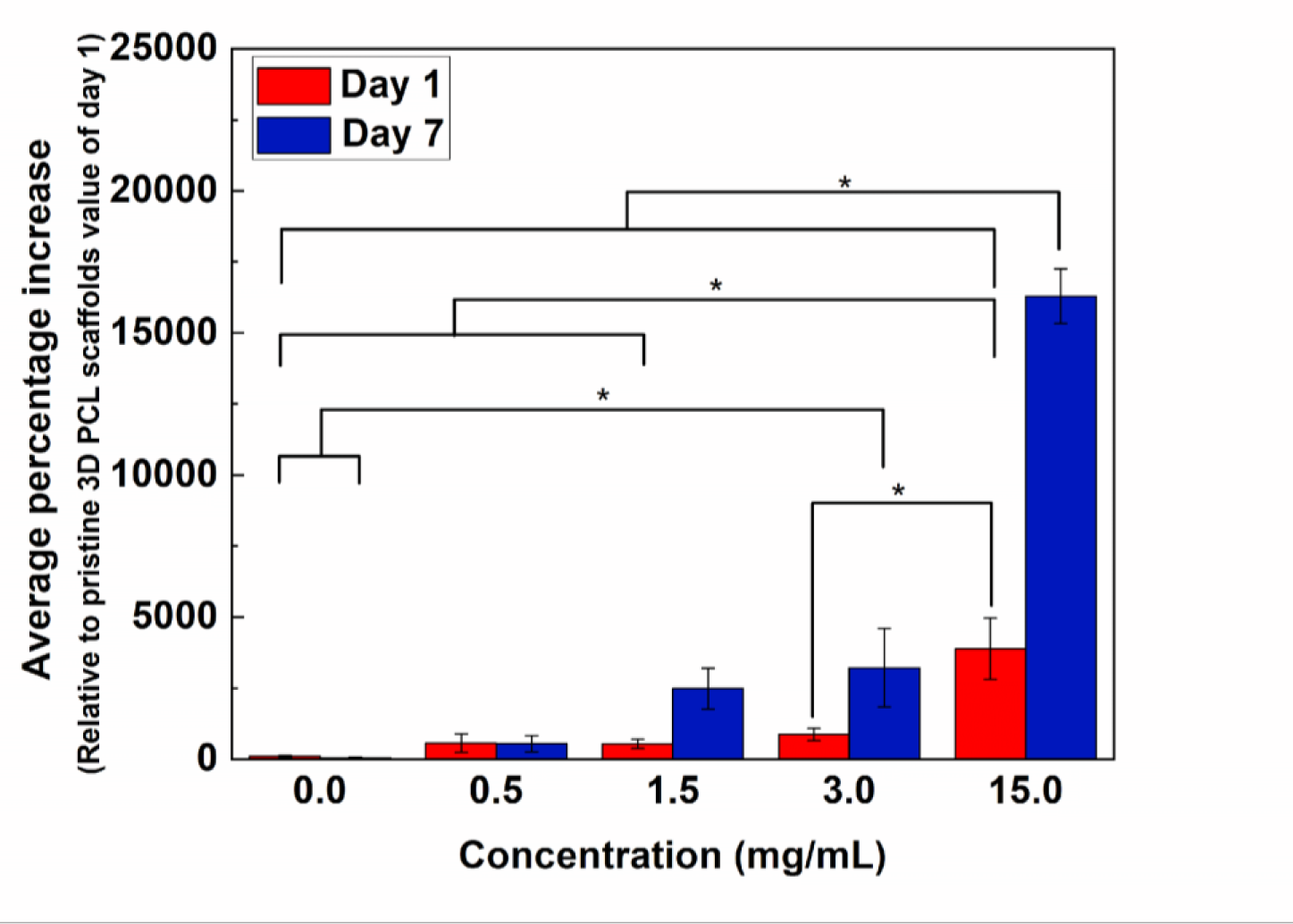
Cell proliferation on pristine 3D PCL scaffolds and Ti_3_C_2_T_x_/PCL scaffolds coated with various Ti_3_C_2_T_x_ MXene concentrations.

### 2.4 Photothermal Effect in Ti_3_C_2_T_x_ MXene and Modulating Neuronal Activity with 3D Ti_3_C_2_T_x_/PCL Scaffold

Ti_3_C_2_T_x_ MXene can convert light to heat efficiently [47], which is critical to trigger the transient capacitive current across the neuronal cell membrane [19]. First, we obtained a calibration curve of the resistance change vs. temperature of a MXene film on a glass slide. The natural logarithm of the relative resistance change (ln (R_t_/R_0_)) was found to be linearly correlated to the reciprocal of temperature (1/T) (ln (R_t_/R_0_) = 1.98×10^3^ 1/T – 6.64; R^2^ = 0.994; **Supplementary Fig. S6**), following Arrhenius law. Next, we measured the resistance of this Ti_3_C_2_T_x_ MXene sample in the presence of an illumination at a wavelength of 640 nm. An incident intensity of about 4.0 x 10^2^ W/mm^2^ (the incident power was 0.485 mW, the focal laser spot diameter was 1.24 µm) for a duration of 10 s led to a resistance change corresponding to a temperature increase of 3.4 ± 0.7 K in Ti_3_C_2_T_x_ MXene, which is similar to other reported results [24]. The intensity is similar to that used for stimulated emission depletion (STED) microscopy [48, 49]. Despite the high intensity, the incident light did not appear to have damaged the stimulated cells, which may be due to the fact that the laser spot was scanned rather than held stationary[50].

Finally, we applied the same incident intensity onto a Ti_3_C_2_T_x_/PCL scaffold with SH-SY5Y cells. To investigate whether photostimulation can cause neural activity, the cells were loaded with Ca^2+^ dye for imaging Ca^2+^ transients. Ca^2+^ are a second messenger of intracellular signal processing and its transient dynamics reflect cellular activity [24, 51]. Since the thermal conductivity of PCL is lower than silicate glass (0.2 W/m·K [52] vs. 1.27 W/m·K [53]), the induced temperature increase of the Ti_3_C_2_T_x_/PCL scaffold could be slightly higher than 3.4 K. As shown in **Fig. 8a-b** and **Supplementary Movie S4**, 640 nm laser light induced Ca^2+^ influx of cell on the Ti_3_C_2_T_x_/PCL scaffold after illumination of about 17 seconds with an intensity of 4.0 x 10^2^ W/mm^2^, while no discernible cellular activity was observed in cells situated on the pristine PCL scaffold (**Supplementary Fig. S5a-b and Movie S5**). These findings show that the MXene coating was required for the photostimulation of neurons on the scaffold, with the temperature increase and optocapacitive effect as the most likely cause. Since an optical beam can be focused to micron and sub-micron diameters, photostimulation can enable spatially precise control of cellular activity within the neuron circuits cultured on the scaffold.

**Fig. 8:**
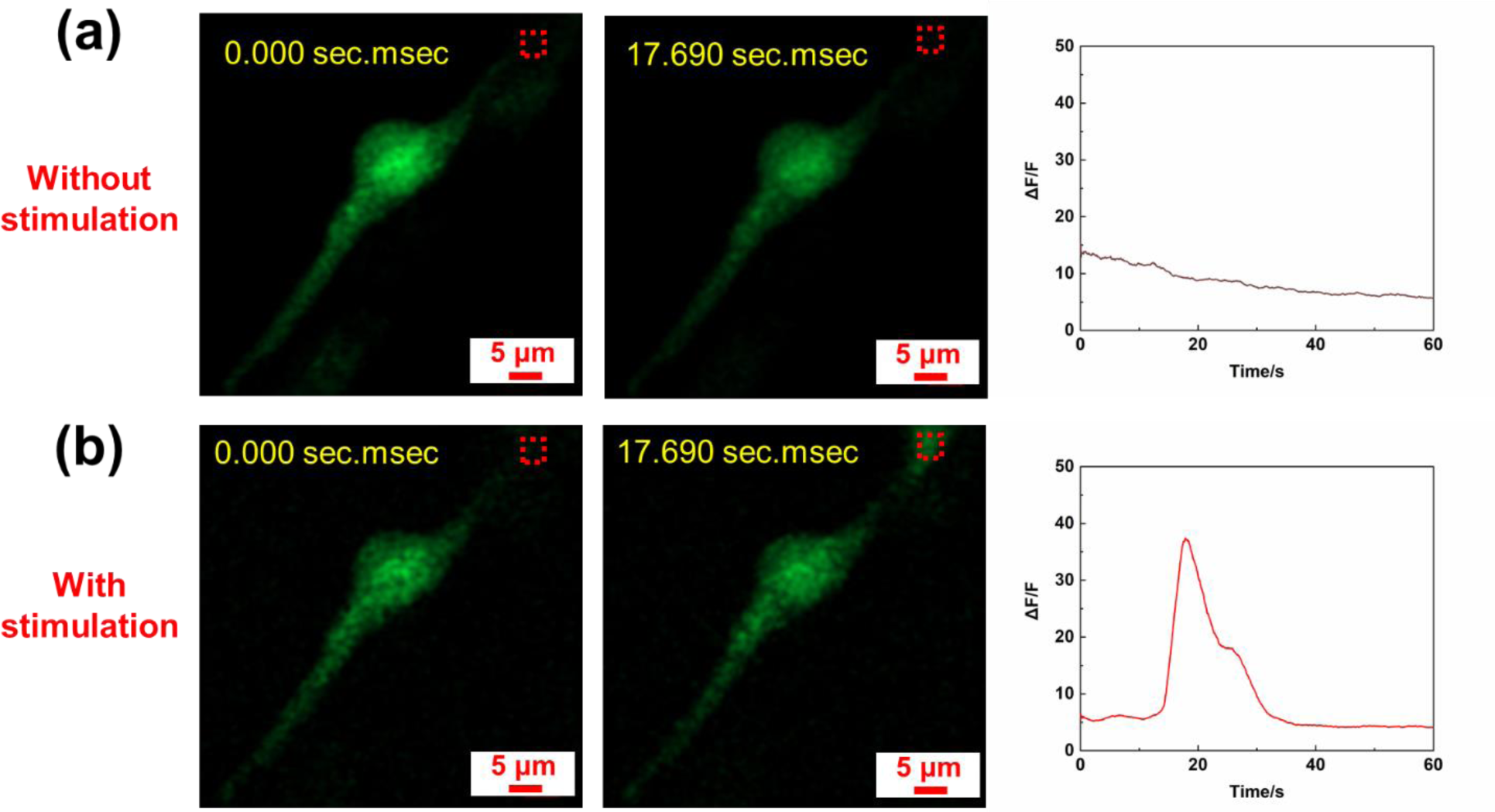
Optical modulation of neuronal cell activity on the Ti_3_C_2_T_x_/PCL scaffold. (a) Fluorescence image of typical SH-SY5Y cells loaded with calcium ions (Ca^2+^) indicator at different time points without 640 nm laser stimulation, followed by a corresponding graph showing the relative change in fluorescence intensity of the region of interest (ROI; red dashed square). (b) Fluorescence image of a typical SH-SY5Y cell at different time points with 640 nm laser stimulation, followed by a corresponding graph showing the relative change in fluorescence intensity of the ROI (red dashed square). Stimulation was initiated simultaneously with the imaging process.

The 3D printed Ti_3_C_2_T_x_/PCL scaffolds can be a versatile tool for neuronal circuit engineering, remote cellular modulation, and neural network computation. **Supplementary Fig. S7** highlights two approaches for 3D neuronal stimulation that will be explored in subsequent investigations. In the first case (**Supplementary Fig. S7a**), neurons are cultivated in conjunction with the 3D printed Ti_3_C_2_T_x_/PCL scaffolds, facilitating the formation of intricate and functional networks. To achieve 3D multi-cellular stimulation, optical beam patterns can be formed using a digital mirror device (DMD) or spatial light modulator. In the second scenario (**Supplementary Fig. S7b**), Ti_3_C_2_T_x_ MXene is patterned in 3D, for example, inkjet printing. This approach enables the controlled deposition of the photothermal material, which can be illuminated to cause cellular stimulation using a collimated optical beam. Furthermore, cell-containing Ti_3_C_2_T_x_ MXene ink, akin to previously reported cell-containing graphene ink [54], can be explored to achieve one-shot assembly of cell-containing 3D scaffolds. Future work will focus on elucidating the underlying molecular mechanisms and the propagation of neural activity throughout the engineered circuit.

## 3. Conclusion

In summary, we have presented an effective approach toward active biointerfaces using Ti_3_C_2_T_x_ MXene coated 3D printed scaffolds for facilitating functional tissue engineering and modulation of neural circuits. Neuron growth was guided by the 3D structure of the scaffold, while the Ti_3_C_2_T_x_ MXene coating increased the hydrophilicity, cell adhesion, and electrical conductivity of the PCL scaffolds. The optocapacitance-based stimulation was achieved by inducing localized temperature changes in the scaffold of several degrees with red light irradiation, and the photoinduced neural activity was confirmed by Ca^2+^ imaging. This work demonstrates an architecture that combines advanced 3D printing techniques with the application of Ti_3_C_2_T_x_ MXene coatings for spatially controlling neuron growth and photostimulation of neural activity. Using holographic light patterns for 2D and 3D photostimulation in combination with scaffolds with more complex structures opens new possibilities for the precise control of neural circuits both *in vitro* and *in vivo*.

## 4. Experimental Section

### 3D MEW based printing of PCL

Scaffolds with microfibers were 3D printed with polycaprolactone (PCL, average Mn 45,000, Merck, Germany) loaded in a 3 mL metallic cartridge at 65 °C with an air pressure of 70 kPa, a speed of 40 mm/s, 24 G nozzle [1.5 mm from the collecting glass slide (VWR, Germany) substrate], voltage of 5.0 kV (3D Discovery Evolution, RegenHU, Switzerland). 3D rectangular models (35 mm × 15 mm) with different SDs (200 μm, 100 μm and 50 μm) and 6 layers were designed by BioCAD (RegenHU, Switzerland) for subsequent 3D printing. PCL scaffolds with random patterns were printed with 50 μm SD, voltage of 5.5 kV and tip to substrate distance of 3 mm. Molten PCL was extruded from the cartridge and the generated fibers were formed in the applied electric field, with ensuing deposition and solidification on the collecting glass slide substrate. To facilitate further measurements and applications, the 3D printed PCL scaffolds were cut into dimensions of 5 mm × 15 mm using a surgical blade (VWR, Germany) and detached from the glass substrate by 70% EtOH (PanReac AppliChem ITW Reagents, Germany) deionized water (DI water) solution.

### Synthesis of Ti_3_C_2_T_x_ MXene

Ti_3_C_2_T_x_ MXene was synthesized according to the previously reported method [55]. Ti_3_AlC_2_ (5 g, Carbon-Ukraine Ltd., Ukraine) was mixed with 50 mL ice-cold solution of HCl (12 M, 37 wt. % VWR, Germany), deionized water, and HF (27–29 M, 48–51 wt. %, Alfa Aesar, Germany) with a volumetric ratio of 30:15:5. The mixture in the propylene bottle equipped with PTFE coated magnet bar was stirred at 500 rpm for 1 hour at 0 °C, followed by stirring at 35 °C for 24 hours. The obtained product was then washed by repeated cycles of centrifugation at relative centrifugal force (RCF) of 2493 xg, 10 °C for 10 min, followed by decanting the acidic supernatant and adding fresh deionized water and mixing, until the pH of the supernatant was neutral.

For delamination, the obtained multilayer Ti_3_C_2_T_x_ MXene particles from the previous step were reacted with 100 mL aqueous solution of LiCl (1.18 M, 99% Sigma Aldrich, Germany) for 24 hours at room temperature (RT, 25°C), while being stirred at 500 rpm. The work-up of the reaction mixture started by centrifugation in the same way as explained above until a very dark black supernatant, containing the delaminated MXene flakes, was formed (usually after 4 times of washing/centrifugation). From this point onward, the supernatants were collected after each centrifugation until the supernatant was no longer dark black and turned into an almost transparent green solution. The collected supernatants were combined and concentrated by high-speed centrifugation at RCF 9420 g, 10 °C, 20 min. The obtained precipitates (containing the MXene flakes) were mixed with 200 mL of deionized water to produce a concentration of 15.0 mg/mL, as measured by gravimetric vacuum filtration.

### Coating of Ti_3_C_2_T_x_ MXene

Ti_3_C_2_T_x_ MXene aqueous dispersion was diluted with EtOH to obtain Ti_3_C_2_T_x_ MXene 70% EtOH dispersion for coating to achieve improved wetting property for the pristine PCL scaffolds. 3.0 mg/mL, 1.5 mg/mL and 0.5 mg/mL Ti_3_C_2_T_x_ MXene 70% EtOH dispersion were prepared by diluting the 15.0 mg/mL Ti_3_C_2_T_x_ MXene water dispersion through adding specific amounts of ethanol, as shown in **Table 1**. DI water was used in all the experiments and all the obtained dispersions were ultrasonicated (Sonorex digitec; Bandelin, Germany) for 30 min to assist the Ti_3_C_2_T_x_ MXene uniform dispersion. For 15.0 mg/mL Ti_3_C_2_T_x_ MXene aqueous dispersion coating, the substrate was detached from the substrate with 10 uL 70% to EtOH and the extra solvent was removed by a Kimwipe tissue paper (Kimberly-Clark, USA), followed by dropping 10 uL 15.0 mg/mL Ti_3_C_2_T_x_ MXene aqueous dispersion with subsequent overturning several times by tweezers, similar to previous reported method [40]. For EtOH containing dispersion, 10 μL Ti_3_C_2_T_x_ MXene 70% EtOH was deposited onto the PCL scaffold and the dispersion spread automatically, only needing tweezers to assist uniform wetting. The coated samples were dried on a hotplate (VWR, Germany) at 50 °C for 2 min in a fume hood (Waldner, Germany).

**Table 1.** Preparation of Ti_3_C_2_T_x_ MXene with different concentrations.

| Ingredient | 3.0 mg/mL | 1.5 mg/mL | 0.5 mg/mL |
| --- | --- | --- | --- |
| 15.0 mg/mL MXene | 200 $\mu\text{L}$ | 100 $\mu\text{L}$ | 33 $\mu\text{L}$ |
| Ethanol | 700 $\mu\text{L}$ | 700 $\mu\text{L}$ | 700 $\mu\text{L}$ |
| Water | 100 $\mu\text{L}$ | 200 $\mu\text{L}$ | 267 $\mu\text{L}$ |

### Contact angle measurement

The wettability of the structures was determined by using the sessile drop measurement on an optical tensiometer (Biolin Scientific Theta Lite, Finland). A 4 µL DI water or 70% EtOH droplet was placed on the tested structures placed on the measuring stage and imaged with a high-speed camera for 10 seconds at an imaging speed of 51 frames per second. The results were analyzed with the Young-Laplace method.

### Electrical resistance measurement

Sheet resistance of the Ti_3_C_2_T_x_ MXene coated scaffolds were measured for evaluation of coating quality according to previously reported method [30]. Two copper electrodes with a length of L were placed parallelly on the rectangular Ti_3_C_2_T_x_ MXene coated PCL scaffolds (length: 15 mm and width: 5 mm) with a distance of D. A digital multimeter (Agilent 34401A, USA) was used for the measurement of resistance [R (Ω)] between the two copper electrodes and the sheet resistance of MXene coated PCL scaffold [ρ (Ω/)] was defined as:

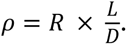

### Scanning electron microscopy

Morphology of MAX phase Ti_3_AlC_2_ and Ti_3_C_2_T_x_ MXene was imaged on a field emission SEM (FESEM, Zeiss Gemini 500) with the InLens detector. Max phase powder particles and etched MXene particles were directly imaged on a conductive carbon tape. Ti_3_C_2_T_x_ MXene was prepared by the following procedures. A diluted aqueous dispersion (0.1 mg/mL) of delaminated Ti_3_C_2_T_x_ MXene flakes was spin-coated on a diced silicon wafer substrate. After drying in the vacuum oven at RT, 10 mBar for 8 h, the substrate was glued to the SEM stub with conductive silver lacquer.

Morphology of 3D PCL scaffolds and 3D Ti_3_C_2_T_x_/PCL scaffolds were characterized with a tabletop SEM (Hitachi TM4000Plus, Japan). For cell containing samples, a fixation of the samples was first carried out in paraformaldehyde fixative solution (PFS; Alfa Aesar, USA) for 30 min with subsequent washing in phosphate buffered saline (PBS; Thermal Fisher, Germany) solution for 3 times, and then the characterization by the SEM was assisted with an ultra coolstage at -30 °C (Deben, UK).

### X-ray diffraction

*Ex situ* powder X-ray diffraction (XRD) pattern of MAX Phase Ti_3_AlC_2_ powder was obtained on an X-ray diffractometer (Aeris Research Edition, Malvern Panalytical Company) using Cu-Kα radiation (λ = 0.15418 nm) at 40 kV and 15 mA at RT in reflection geometry.

### Culture and differentiation of human SH-SY5Y cells

For sterilization of samples before cell culture, 3D printed PCL and 3-time coated Ti_3_C_2_T_x_/PCL scaffolds were immersed in 70% EtOH solution for 2 h, and dried under UV illumination in a biological safety cabinet (BSC; Kojair, Finland) for 1 h. The scaffolds were incubated in culture medium (CM) overnight before cell seeding and kept in a humidified cell culture incubator (5% CO_2_ and 37 °C; Binder, Germany). SH-SY5Y cells were seeded on both pristine 3D PCL and Ti_3_C_2_T_x_/PCL scaffolds with a cell density of 3 × 10^5^ cells/cm^2^ with medium refreshed every two days. The CM contained Dulbecco’s Modified Eagle’s Medium/Nutrient Mixture F-12 (DMEM/F-12; HyClone, USA), 15% (v/v) heat inactivated newborn bovine calf serum (HyClone, USA), 1% (v/v) penicillin-streptomycin (10,000 U/mL; ThermoFisher Scientific, Germany). The differentiation medium (DM) was prepared by supplementing 10 μM retinoic acid (RA; Merck, Germany) in the CM. **Viability staining:** For cell viability, distribution and proliferation evaluation, 8 μg/mL Calcein AM (Merck, Germany), 4 μg/mL PI (Merck, Germany) and 12 μg/mL Hoechst 33342 (Miltenyi Biotec, Germany) were used to stain the live cells, dead cells and nuclei, respectively. Briefly, samples were incubated with Calcein AM for 15 min in an incubator, following with addition of PI and Hoechst for an additional 15 min incubation in an incubator. After samples washing with fresh CM, images were obtained with a confocal microscope (LSM 900) (Zeiss, Germany) and analyzed with Zen software (Zeiss, Germany). For proliferation analysis, the area of stained live cells was extracted by ImageJ/Fiji software [56].

### Immunofluorescence staining

Samples were washed in PBS solution with subsequent fixation in PFS for 30 min at RT after 2-day culture in CM, before and after 3 days culture in DM. Fixed samples were rinsed 3 times in PBS solution, before being blocked and permeabilized by a PBS solution supplemented with 10% (v/v) horse serum (HS, MP Biomedicals) and 0.3% (v/v) Triton X-100 (Merck, Germany) at RT. After 3 times washing in PBS, samples were incubated in a 10% (v/v) HS PBS solution supplemented with primary mouse anti-β-tubulin III antibody (1:100; Merck) overnight at 4 °C. Thereafter, samples were washed in PBS for 3 times and Alexa Fluor 555 conjugated secondary goat anti-mouse IgG antibody (1:100; Merck, Germany) was added for immunostaining in the dark for 2 hours at RT. F-actin and nuclei were stained by AlexaFluor™488 phalloidin (ThermoFisher Scientific, Germany) and DAPI (Merck, Germany) in the dark for 1 hour at RT, respectively. After samples washing in PBS, ProLong™glass antifade mountant (ThermoFisher Scientific, Germany) was used to mount the samples on a glass slide before imaging. Images were obtained with Zeiss LSM 900 and analyzed with Zeiss Zen software.

### Photothermal effect studies

Material electrical resistance changes with temperature by following Arrhenius relationship [24]

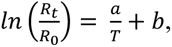

where R_t_ is the measured electrical resistance at temperature T, R_0_ is the measured electrical resistance at temperature 303.15 K (30 °C), a is the slope of the curve and b is the curve intercept. The MXene electrode was prepared by adding 10 µL10 mg/mL MXene water solution on the glass slide with a 2 mm trench made by two parallel scotch tape (3M, USA) and dried on a 50 °C hot plate (VWR, Germany) for 5 min in the fume hood. The electrode temperature was controlled by submerging in 70 °C PBS solution with a P4010 thermal meter (Dostmann electronic, Germany) for temperature monitoring and a 34401A digit multimeter (Agilent, USA) for electrical resistance measurement. For the photothermal effect measurement of the MXene sample, the fluorescent latex beads (2 µm, fluorescent red; Merck, Germany) were diluted by 1000 times and added in the MXene electrode to assist in finding the focal plane when applying photo stimulation. Confocal microscope (LSM 900) (Zeiss, Germany) was used for applying laser stimulation and the laser power was measured by an 843-R power meter (Newport, USA). The laser focus spot size was estimated with the following equation:

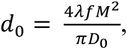

where *d*_0_ is the focal laser spot diameter, *λ* is the wavelength, *M*^2^ is the beam quality parameter, f is the focal length, and *D*_0_is the input beam diameter. The diameter is 1.24 µm.

### Calcium imaging

Samples were loaded with 5 μM Fluo-4 AM (ThermoFisher Scientific, Germany) in CM for 30 min in a cell culture incubator (37 °C and humidified 5% CO_2_), while 0.02% (w/v) Plu was added to assist the calcium dye dispersion in CM as per the manufacturer’s instruction. Thereafter, samples were washed in fresh CM and imaged on a confocal microscope (Zeiss LSM 900) with a mounted incubator (XLmulti S2 DARK, Germany). The 488 nm and 640 nm lasers were turned on during the imaging process, with a line scan frequency of ∼3 KHz and a pixel dwell time of ∼2 µs, using bidirectional scanning. The pixel size is 397 nm for the cell on the MXene coated scaffold imaging, while it is 248 nm for the cell on the pristine scaffold. ImageJ/Fiji software was used for acquired image and movie processing. Background fluorescence was subtracted prior to fluorescence change analysis of ROIs.

### Statistical analysis

All the obtained quantitative data were demonstrated as mean ± standard deviation and measurements were executed in triplicate. Two-way analysis of variance (ANOVA, Bonferroni post hoc test) was performed with a significance level set of < 0.05 if homogeneity of variance (Brown–Forsythe test) was satisfied (> 0.05). Otherwise, the significance level was set to < 0.01. All the statistical analyses were carried out by using OriginPro 2019 software (OriginLab, USA).

## Supporting information

MEW based 3D printing of PCL scaffold with a SD of 50 micron.

Contact angle measurement with 70% EtOH droplet.

Contact angle measurements with DI water droplets on both 3D printed pristine PCL and MXene coated PCL scaffolds.

Optical modulation of neuronal cell activity on the Ti3C2Tx/PCL scaffold without and with 640 nm laser stimulation.

Optical modulation of neuronal cell activity on the pristine PCL scaffold without and with 640 nm laser stimulation.

## 6. Acknowledgements

Financial support of the Max Planck Society is gratefully acknowledged. The authors thank Dr. Davood Sabaghi for support with the XRD measurements. JL would like to acknowledge Mr. Fu-Der Chen, Dr. Andrei Stalmashonak and Mr. Frank Weiß for technical support.

