## Supplementary information for "3D Printed Ti_3_C_2_T_x_ MXene/PCL Scaffolds for Guided Neuronal Growth and Photothermal Stimulation"

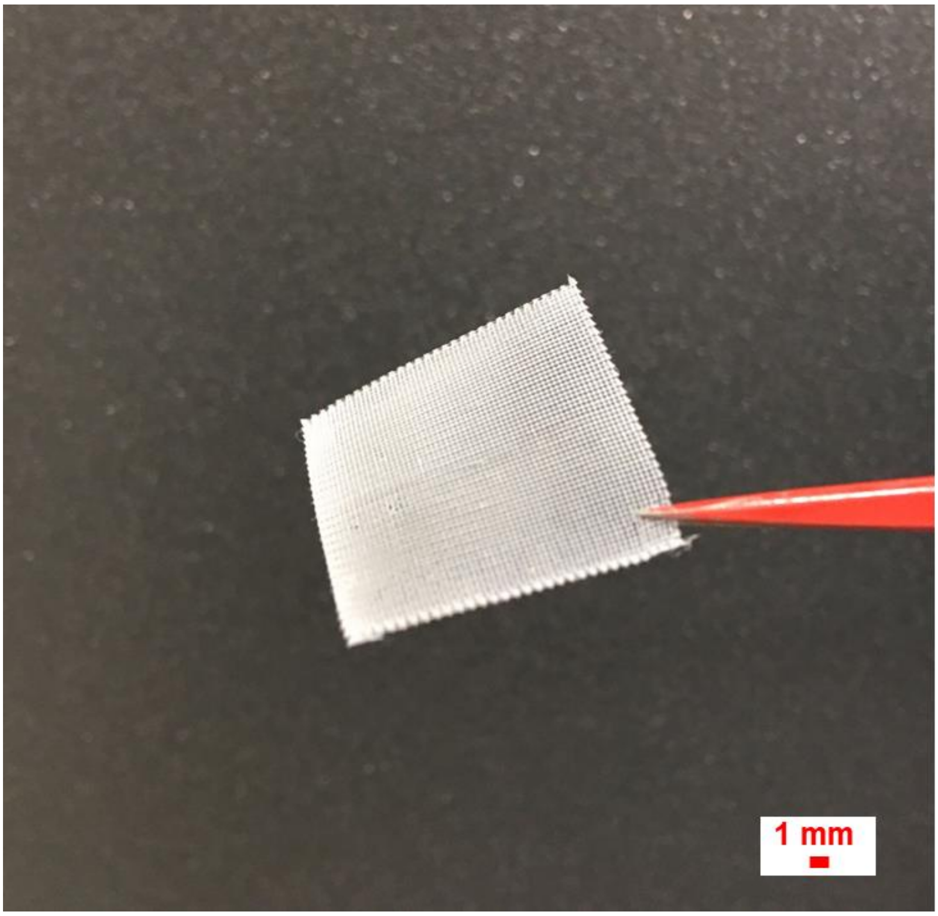


**Supplementary Fig. 1 | MEW based 3D printed PCL scaffold could be mechanically stable.** 3D printed PCL scaffold with 90 layers and a strand distance (SD) of 300 μm is handled with tweezers.

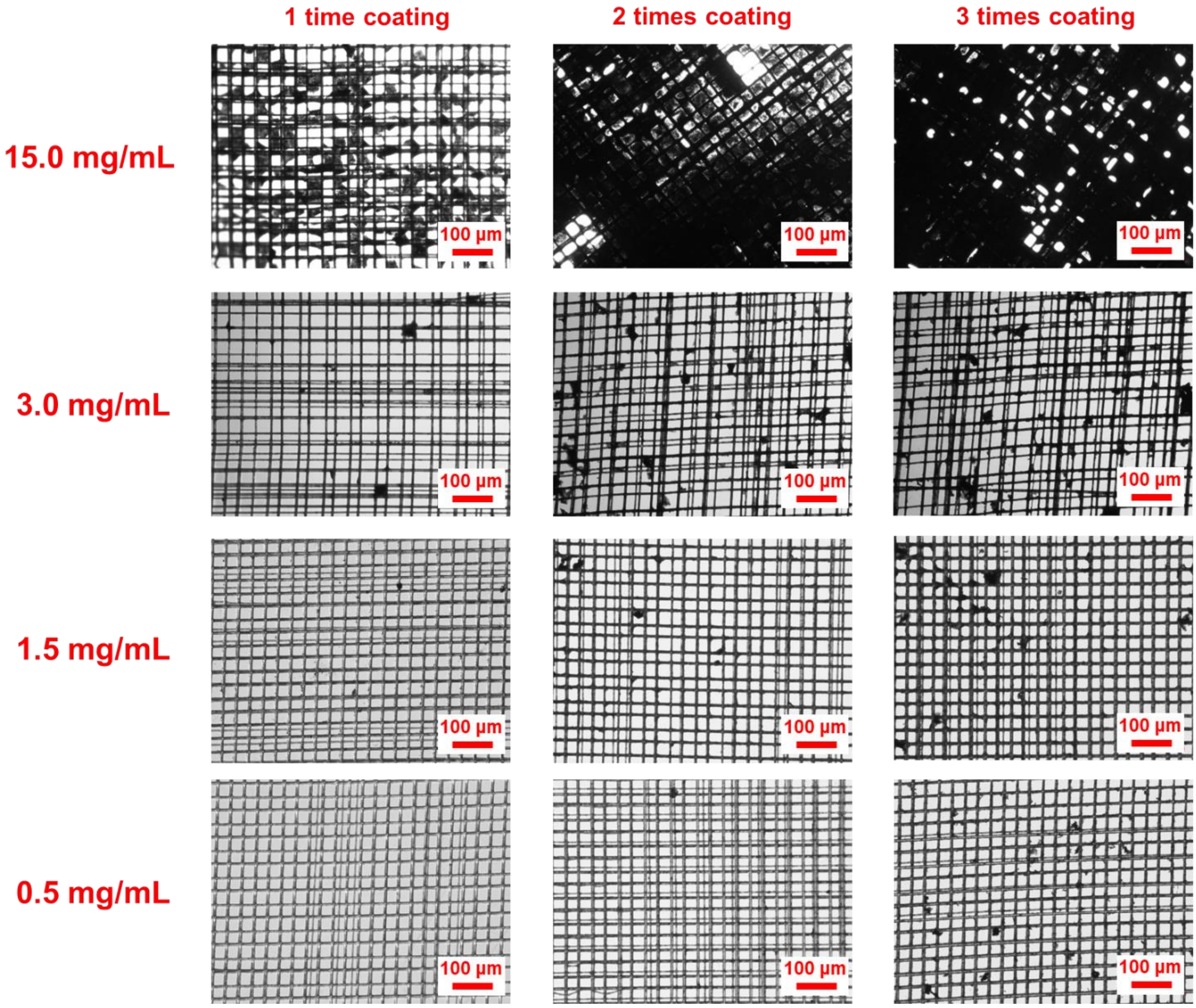


**Supplementary Fig. 2 | $Ti_3C_2T_x$/PCL scaffold morphology with different $Ti_3C_2T_x$ coating concentration and coating time.** Optical microscopy images for the morphology of $Ti_3C_2T_x$/PCL scaffolds (SD ranges from 1 to 50 μm) following different times of coating (1-3) with 15.0 mg/mL, 3.0 mg/mL, 1.5 mg/mL and 0.5 mg/mL $Ti_3C_2T_x$ MXene dispersions, correspondingly.

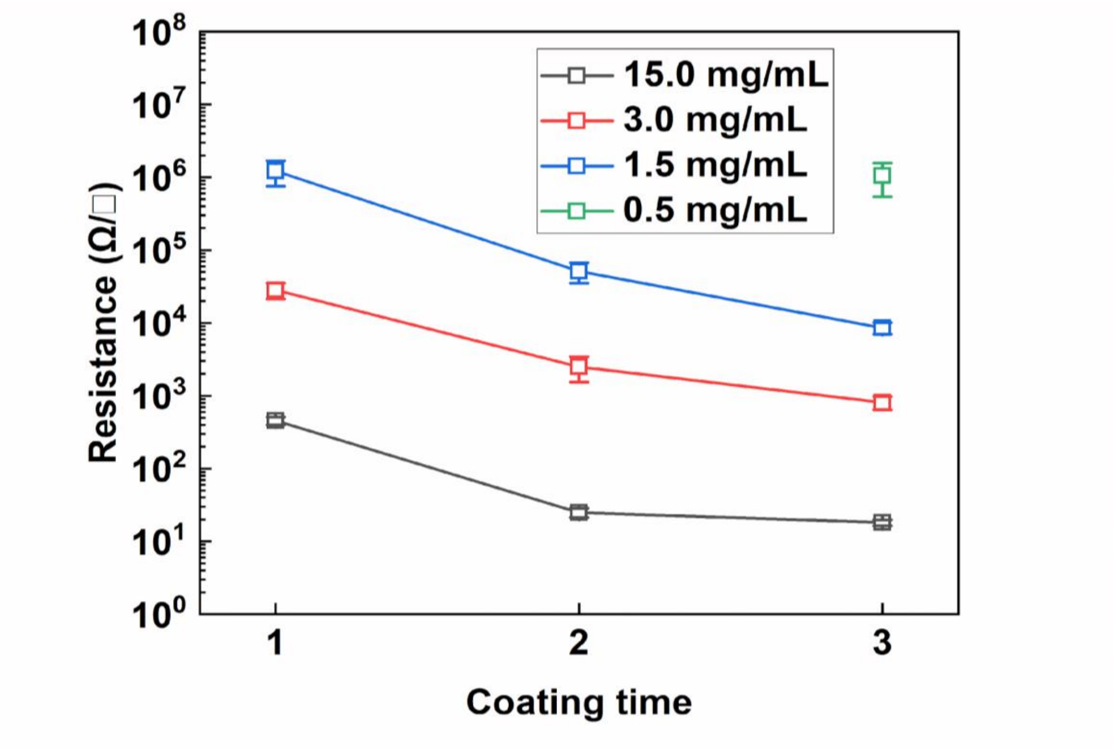


**Supplementary Fig. 3 |** Sheet resistance of $Ti_3C_2T_x$ MXene coated PCL scaffolds with different concentrations of $Ti_3C_2T_x$ MXene dispersion and coating times.

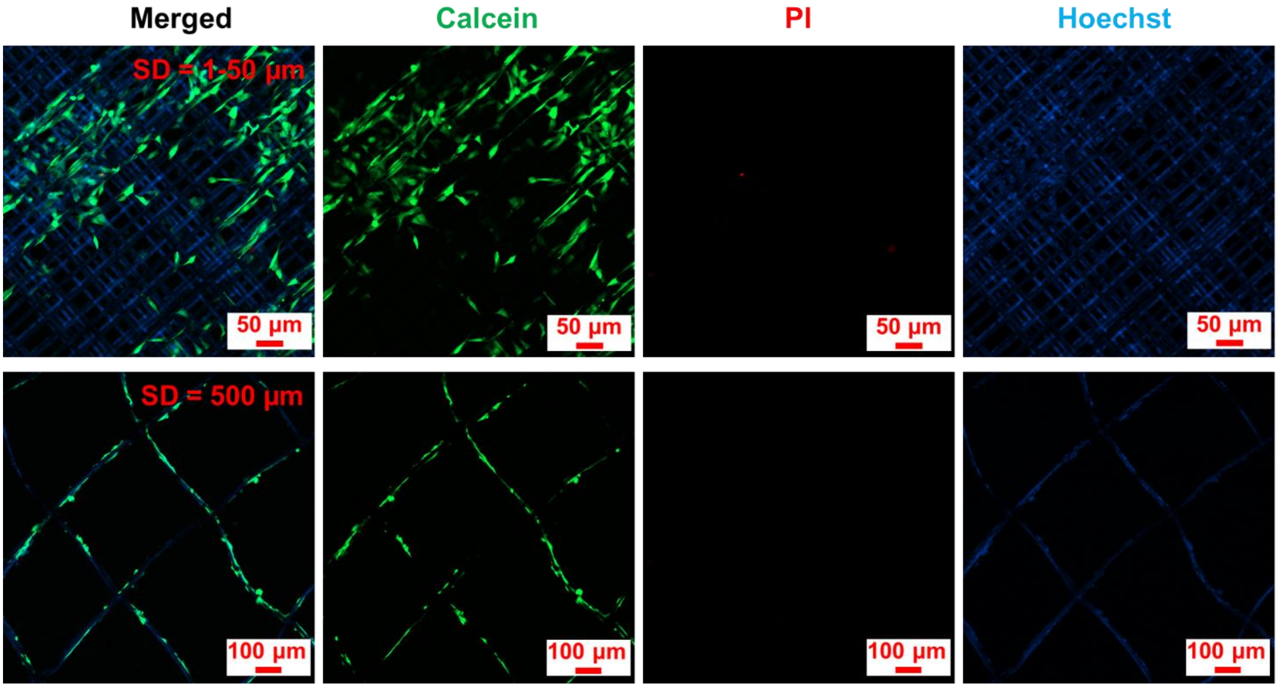


**Supplementary Fig. 4 | Cell viability and morphology on the 3D MXene/PCL scaffolds with different SDs (1-50 μm and 500 μm) after 6 days culture.** Live cell (Calcein; green), dead cell [propidium iodide (PI); red] and cell nucleus with PCL substrate (Hoechst; blue).

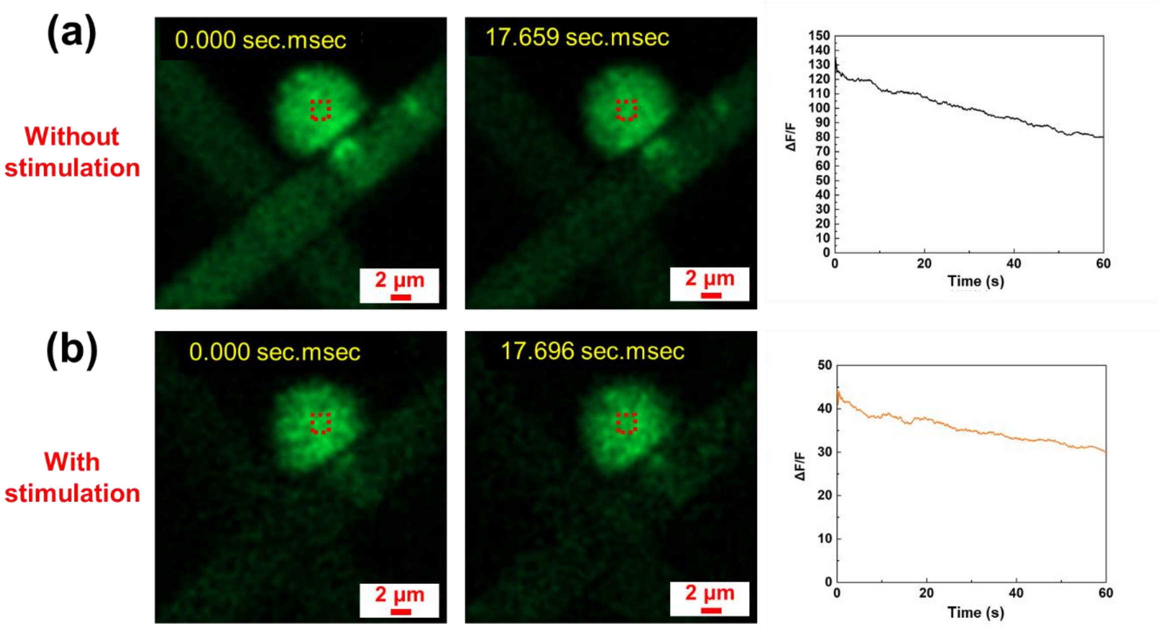


**Supplementary Fig. 5 | Optical modulation of neuronal cell activity on the pristine PCL scaffold.** (a) Fluorescence image of typical SH-SY5Y cells loaded with $Ca^{2+}$ indicator at different time points without 640 nm laser stimulation, followed by a graph showing the relative fluorescence intensity change of the ROI (red dashed square). (b) Fluorescence image of a typical SH-SY5Y cell at different time points with 640 nm laser stimulation, followed by a graph showing the relative fluorescence intensity change of the ROI (red dashed square). Stimulation was initiated simultaneously with the imaging process.

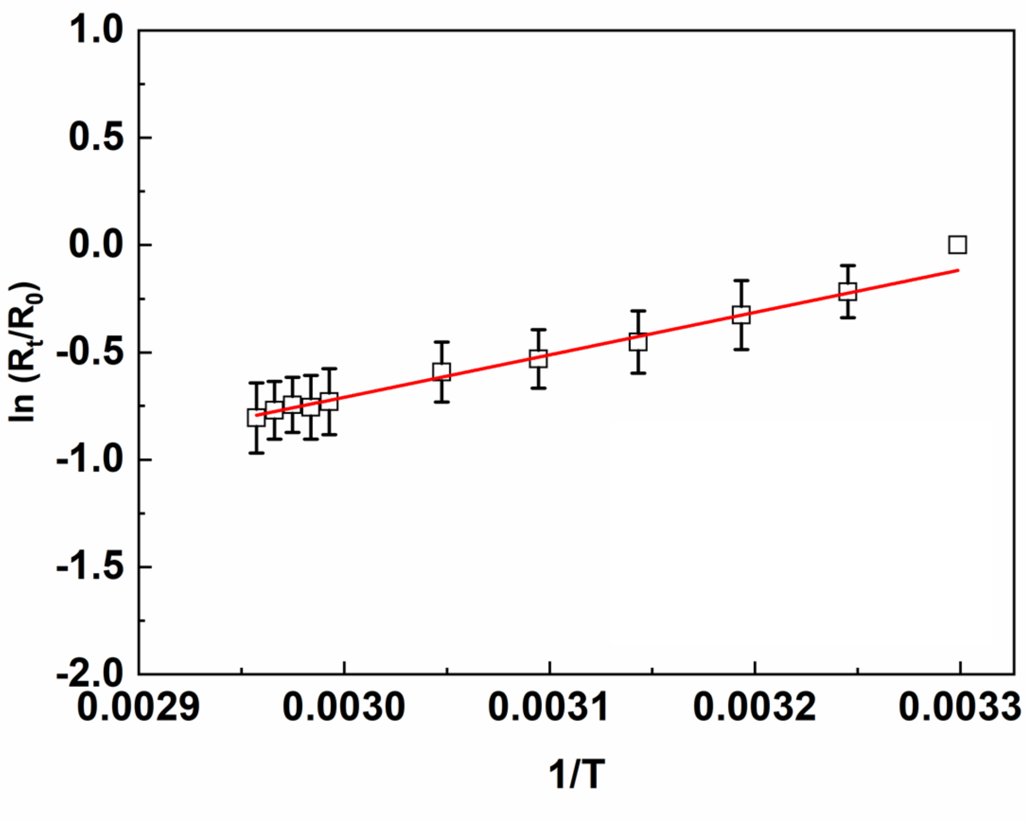


**Supplementary Fig. 6 | Temperature-dependent electrical resistance of $Ti_3C_2T_x$ MXene and photothermal effect in the 3D $Ti_3C_2T_x$/PCL scaffold.** The natural logarithm of the relative resistance change (ln ($R_t/R_0$)) of $Ti_3C_2T_x$ MXene as a function of reciprocal of temperature (1/T), where $R_t$ is the measured $Ti_3C_2T_x$ MXene electrical resistance, $R_0$ is the $Ti_3C_2T_x$ MXene electrical resistance at 30°C and T is the temperature in Kelvin. Measurement was taken from 30°C to 65°C.

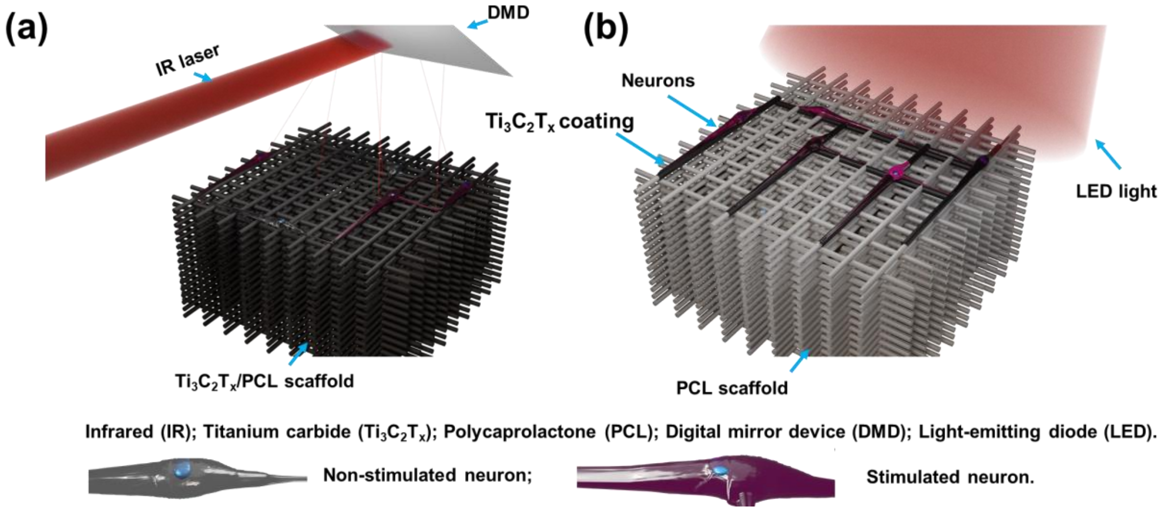


**Supplementary Fig. 7 | Two proposed approaches for future work.** (a) Addressable stimulation on the neural circuits formed on the $Ti_3C_2T_x$/PCL scaffold using a programmable light pattern enabled by DMD. (b) Addressable stimulation of the neural circuits using patterns of $Ti_3C_2T_x$.

**Supplementary Movie 1**. MEW based 3D printing of PCL scaffold with a SD of 50 μm.

**Supplementary Movie 2**. Contact angle measurement with 70% EtOH droplet.

**Supplementary Movie 3**. Contact angle measurements with DI water droplets on both 3D printed pristine PCL and MXene coated PCL scaffolds.

**Supplementary Movie 4**. Optical modulation of neuronal cell activity on the $Ti_3C_2T_x$/PCL scaffold without and with 640 nm laser stimulation.

**Supplementary Movie 5**. Optical modulation of neuronal cell activity on the pristine PCL scaffold without and with 640 nm laser stimulation.
